# Vritra: gene-specific reference construction for species-resolved functional profiling of metagenomic and metatranscriptomic data

**DOI:** 10.64898/2026.09.17.752478

**Authors:** Boyan Zhou, Menghan Liu, Gary Curhan, Gregory Peck, Hyon Choi, Lama Nazzal, Huilin Li

**Affiliations:** Division of Biostatistics, Department of Population Health, New York University School of Medicine, New York, New York 10016, USA; Department of Biological Sciences, Columbia University in the City of New York, New York, NY, 10027, USA; Department of Medicine, Brigham and Women’s Hospital, Harvard Medical School, Boston, MA 02115, USA; Department of Surgery, Robert Wood Johnson Medical School, Rutgers University, New Brunswick, NJ 08901, USA; Division of Rheumatology, Allergy and Immunology, Department of Medicine, Massachusetts General Hospital, Harvard Medical School, Boston, MA 02115, USA; Department of Medicine, Division of Nephrology, NYU Langone Medical Center, New York, NY, 10016, USA

**Keywords:** metagenomics, metatranscriptomics, microbial functional genes, species-level attribution, gene-specific reference databases

## Abstract

Species-resolved profiling of microbial functional genes is important for linking microbial community composition to biological function, but existing functional profiling approaches are not designed to systematically define user-specified genes and resolve their contributing species. We developed Vritra (**V**ersatile gene-guided **R**eads-identification with **I**mpartial **T**axonomic **R**efinement and **A**ssignment), a framework for constructing gene-specific reference databases for metagenomic and metatranscriptomic data. Vritra expands and refines the UniRef-based sequence space for user-specified target genes using sequence-similarity network connectivity and functional annotations, retains related homologs as decoys to reduce assignment ambiguity, and links the resulting sequences to standardized microbial taxonomy. Across genes involved in oxalate, urate, and bile acid metabolism, Vritra recovered >97% of sequences represented by established annotation resources for most evaluated genes while substantially expanding the represented sequence space. In real microbiome datasets, the expanded references increased recovery of target-gene reads by up to threefold for poorly annotated genes while maintaining high sequence identity. Application to population-based and publicly available microbiome datasets enabled species-resolved profiling of functional genes and revealed gene-specific associations with microbial taxa, dietary factors, and disease-related phenotypes. Vritra provides a scalable framework for translating continuously expanding sequence resources into gene-specific, species-resolved references for microbiome studies.

**Importance:** Microbial genes encode functions that can influence human health, yet it is often difficult to determine which microbial species carry out specific functions in complex microbiome communities. Existing approaches are well suited for broad functional profiling but provide limited flexibility for studying individual genes, particularly when those genes are poorly characterized or represented inconsistently across reference resources. Vritra addresses this gap by providing a framework for building gene-specific reference databases that connect functional genes with the microbial species that carry them. The approach can be applied to both metagenomic and metatranscriptomic data and can accommodate user-specified target genes, including genes from emerging or incompletely characterized pathways. By enabling species-resolved profiling of selected microbial functions, Vritra can help researchers investigate how specific microbial activities vary across individuals and relate to dietary, ecological, and disease-related factors. This framework provides a practical approach for studying microbial functions that may be overlooked by broad functional profiling methods.

## Introduction

Taxonomic classification of metagenomic (MGX) and metatranscriptomic (MTX) data is fundamental to microbiome research, yet many investigations now extend beyond community composition to interrogate specific microbial functions. As studies increasingly seek to link microbial activity to host physiology and disease, there is growing need for methods that can sensitively detect individual microbial genes or curated gene sets with defined biological roles—and accurately attribute them to their encoding species.

Motivated by our long-standing focus on the “oxalobiome”, we recently developed an oxalate-centered analysis pipeline^1^ that resolved species-level contributions to the oxalate-degradation genes *frc* and *oxc*^1^. By constructing a targeted reference database from InterPro^2^ and leveraging curated functional family and domain annotations, this framework enabled high-resolution functional-taxonomic mapping. Its application across large MGX and MTX population cohorts has substantially advanced our understanding of microbial oxalate degradation and its relevance to kidney stone formation^3, 4^.

However, this oxalate-focused approach could not be generalized to other microbial gene sets of emerging biomedical interest for which clearly defined sequence sets are unavailable in InterPro. For examples, two recently published pathways—bile acid 7α-dehydroxylation pathway^5, 6^ and high-capacity urate metabolism pathways^7, 8^— comprise multi-gene systems with demonstrated or suspected roles in gallstone disease and hyperuricemia, respectively. Each pathway consists of a few recently discovered or under-characterized genes—and identifying their homologs and taxonomic origins from MGX/MTX data remains challenging, because existing reference databases lack curated functional representations and clear mapping to corresponding gene names^9, 10, 11^.

Although tools such as HUMAnN excel at broad functional profiling of microbial communities, they are not designed to systematically link specific functional genes to their contributing species^12, 13^. HUMAnN primarily quantifies gene families using UniRef90, which groups protein sequences sharing ≥90% sequence identity^14^. However, a single UniRef90 cluster can encompass sequences from multiple microbial species^14^ (Fig. 1a, Step 3), thereby limiting species-level attribution of individual functional genes. The development of the Genome Taxonomy Database (GTDB)^15^, which provides a standardized microbial taxonomy and defines species boundaries at approximately 96% average nucleotide identity (ANI)^16^, provides an opportunity to establish a consistent taxonomic framework for species-resolved functional analysis.

**Fig. 1.**
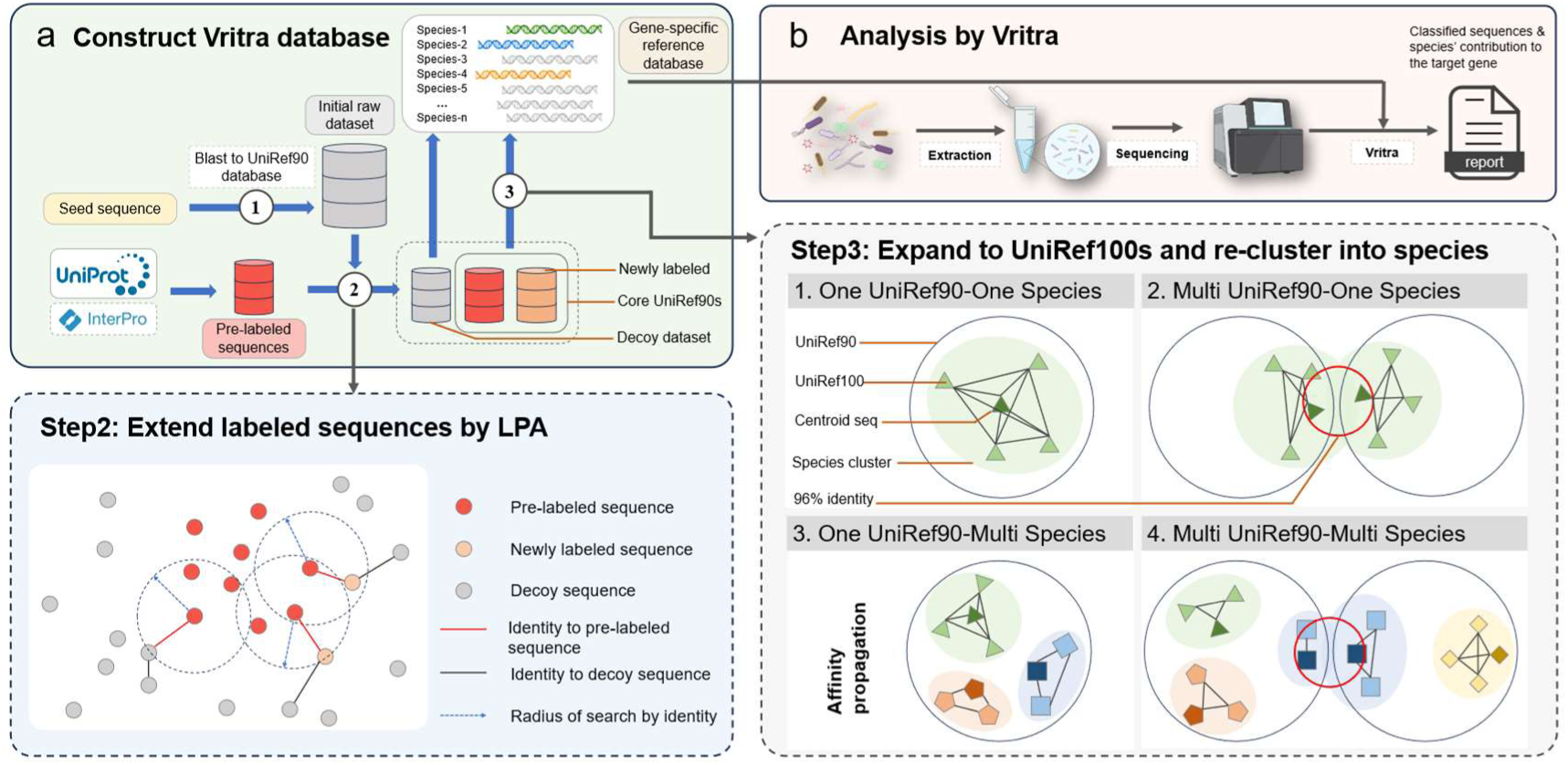
Workflow of Vritra. **a** Construction of Vritra database: Step1. Searching the “seed sequence” of target gene against the complete UniRef90 database to obtain an “initial raw dataset”. Step 2. An annotated set of protein sequences is retrieved from UniProt or InterPro using the scientific name of the target gene or protein. These “pre-labeled” sequences are expanded into the “core UniRef90s” within the initial raw dataset using a Label Propagation Algorithm (LPA), and the remaining sequences constitute the “decoy dataset”. Step 3. The “core UniRef90s” are mapped to their corresponding UniRef100 entries and re-clustered based on taxonomic relationships and pairwise sequence identity. Representative sequences are selected using either centroid choice or the affinity propagation algorithm. The final “gene-specific reference database” comprises these representative sequences together with the “decoy dataset”. **b** Analysis by Vritra: Raw sequencing reads from real data are aligned to the “gene-specific reference database”, and hits mapping to the “decoy dataset” are removed. The remaining reads are assigned to their best-matching reference sequences, yielding species-resolved abundance tables reported as read counts and RPKM values.

Beyond the challenge of species-level attribution, accurately identifying the UniRef90 clusters corresponding to specific target genes is further complicated by inconsistent protein-name annotations^17^. Because UniRef draws its names from diverse sources (UniProtKB, UniParc, RefSeq^18^, PDB^19^, Ensembl^20^), sequences within a cluster may inherit different naming conventions^21^, based on domain architecture, catalytic activity, or hierarchical function. Consequently, text-based searches alone may incompletely or inaccurately define the sequence space corresponding to a target gene. For example, searches for *xdhA* may fail to capture all relevant xanthine dehydrogenase sequences because subunit organization and protein annotations vary across species^22^. Thus, gene symbols alone do not provide a reliable basis for defining a complete sequence set for a target function.

Comprehensive sequence references such as UniRef100 provide broader sequence coverage but are challenging to use for direct read alignment in large-scale human gut MGX/MTX datasets because of their size (>100 GB)^23^. Reduced or customized reference databases can improve computational tractability^12, 24, 25^, but may sacrifice sequence completeness or introduce inconsistencies in gene definitions and taxonomic assignments. These limitations are particularly consequential for target-gene analysis, where the reference must simultaneously provide comprehensive sequence representation, consistent functional definitions, and taxonomically resolved species assignments.

To address this gap, we introduce Vritra, a framework that integrates gene definitions, sequence retrieval, and standardized microbial taxonomy to construct gene-specific, species-resolved reference databases for MGX and MTX data. Vritra expands and refines the UniRef-based sequence space for user-specified target genes and links the resulting sequences to taxonomically resolved microbial species, enabling reproducible species-level profiling of selected microbial functions while maintaining a computationally tractable reference space.

## Results

### Workflow of Vritra

In this study, we developed Vritra (<u>V</u>ersatile gene-guided <u>R</u>eads-identification with Impartial <u>T</u>axonomic <u>R</u>efinement and <u>A</u>ssignment), a framework for constructing gene-specific reference databases and enabling species-resolved profiling of target genes from MGX/MTX sequencing data. Vritra consists of two main components: (1) construction of a gene-specific, taxonomically resolved reference database (Fig. 1a) and (2) application of this reference to MGX/MTX sequencing data to generate species-resolved gene abundance profiles (Fig. 1b).

#### Sequence-space expansion

Database construction in **Vritra** starts with a user-specified seed protein sequence representing the target gene (Fig. 1a). Vritra searches the seed sequence against the UniRef90 database to retrieve a comprehensive set of homologous sequences (Fig. 1a, “initial raw dataset” in Step 1), ensuring broad coverage of gene variants while keeping the search space well defined. This sequence-based retrieval avoids the ambiguity of text-based annotation searches and provides a systematic starting point for defining the sequence space of the target gene.

#### Functional refinement

To refine the initial dataset and remove spurious sequences, Vritra applies two complementary strategies, depending on the extent of gene annotation, to construct the “core UniRef90 dataset” (Fig. 1a, Step 2). For genes with established annotation, relevant sequences (i.e., “pre-labeled sequences” in Fig. 1a) are retrieved from UniProt (optionally supplemented with InterPro) using the scientific name of the gene or protein. These sequences are then expanded through a Label Propagation Algorithm^26^ (LPA; Fig. 1a, Step 2, Methods) within the initial raw dataset to form the “core UniRef90 dataset”, leveraging sequence-network connectivity to identify additional homologs that may have incomplete, inconsistent or higher-level annotations. For genes with limited annotation, such as those represented by only a few sequences or entirely absent from databases like InterPro or UniProt, Vritra provides an exploratory mode that broadens candidate retrieval beyond curated entries (Methods). Whereas iterative approaches such as PSI-BLAST^27^ can recover increasingly remote homologs and consequently introduce functionally ambiguous matches (e.g., <35% sequence identity)^28, 29^, Vritra uses an identity-based heuristic with adjustable similarity thresholds to restrict candidate sequences to a more functionally relevant range.

Sequences from the initial raw dataset that are not incorporated into the “core UniRef90 dataset” are retained as a “decoy dataset” rather than discarded (Fig. 1a, Step 2). Retaining them as an explicit negative reference allows Vritra to identify and exclude reads that preferentially map to homologous or paralogous proteins within the same protein superfamily, thereby reducing ambiguity in downstream target-gene detection.

#### Taxonomic disambiguation

To establish species-resolved attribution, Vritra next expands sequences in the core UniRef90 dataset to their corresponding UniRef100 sequence space and links these sequences to the standardized taxonomy (Fig. 1a, Step 3). Because UniRef90 clusters do not necessarily correspond uniquely to individual microbial species, Vritra explicitly resolves UniRef90-species relationships into four predefined categories (Fig. 1a, Step 3; Methods). Representative sequences are then selected for each taxonomically resolved species and combined with the decoy dataset to generate the final gene-specific reference database. This step establishes a direct link between the target-gene sequence space and standardized microbial species while avoiding the need to align sequencing reads against the full UniRef100 database.

For downstream profiling, MGX/MTX reads are aligned to the resulting gene-specific reference database using Diamond2^30^ (Fig. 1b). Reads are assigned to the best-matching target-gene representative and reads preferentially mapping to decoy sequences are excluded. Vritra thereby generates species-resolved abundance profiles for the target gene, reported as read counts and RPKM (reads per kilobase of gene length per million mapped reads). The resulting framework connects target-gene definitions, sequence representation and microbial taxonomy within a unified reference.

### Characterization of Vritra-derived gene-specific reference databases

To examine the general applicability of this framework across genes with different levels of annotation, we selected three bacterial gene sets spanning a broad range of annotation availability: an oxalate-degradation set (*frc* and *oxc*)^1^, a urate-degradation set (*ygeX*, *hyuA*, *ygeW*, *ygfK*, *ssnA*, *ygeY*, and *xdhA*)^8^, and two genes (*baiB* and *baiE*)^6^ from the bile-acid-induced (bai) operon. These genes span a gradient from well-annotated (e.g., *frc* and *oxc)* to less-characterized genes (e.g., *xdhA* and *hyuA),* with the remaining genes falling between these extremes. For the oxalate- and urate-degradation genes, we compared Vritra-derived sequence sets with those generated using well-annotated InterPro entries (version 106.0), using the latter as an established reference for assessing recovery. In contrast to the oxalate- and urate-pathway related gene sets, the bai genes lack sufficient InterPro coverage (only 10 sequences are available for *baiB* and none for *baiE*). Therefore, we present their analysis only as an application of Vritra to real data.

#### Recovery and expansion of established sequence space

We first asked whether Vritra could recover sequence representations supported by existing annotation resources while extending coverage beyond those annotations. Because the InterPro-derived sets are not standardized to UniRef clusters, whereas Vritra relies on representative sequences, the two datasets cannot be compared on a one-to-one basis. Therefore, we defined an “overlap” as pairs of sequences sharing >80% alignment coverage and >90% sequence identity between the two methods, a criterion analogous to the clustering principle underlying UniRef90. Using this criterion, Vritra recovered >97% of InterPro-derived sequences for all evaluated genes except *ssnA*, for which the recovery was 92.9% (Fig. 2a). In addition, Vritra substantially expanded sequence representation, increasing the number of sequences by approximately threefold for *xdhA*, *ygeX*, and *hyuA*. These results indicate that Vritra can retain sequence representations supported by established annotations while extending the reference beyond the sequences captured by existing annotation resources, particularly for genes with limited annotation.

**Fig. 2.**
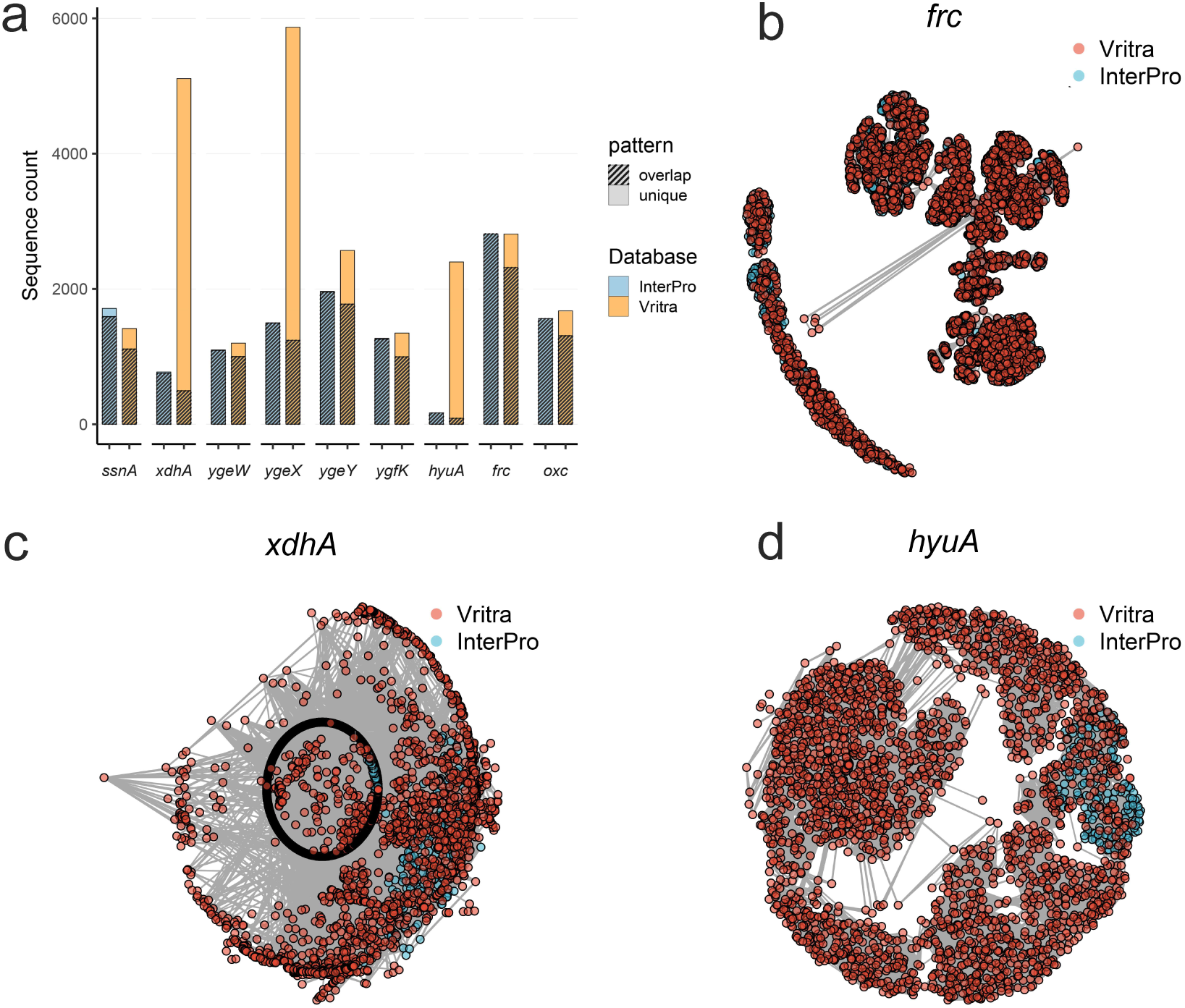
Comparison between Vritra-constructed databases and InterPro databases. **a**, Number of protein sequences in databases generated by Vritra and obtained from InterPro for two gene sets. The shaded region indicates the overlapping sequences between the two databases, defined as sequences sharing >80% coverage and >90% pairwise identity. **b–d**, Sequence-level coherence of sequences obtained by the two methods for three genes: **b**, *frc*; **c**, *xdhA*; **d**, *hyuA*.

#### Sequence-level coherence of the expanded sequence space

We next examined whether the additional sequences identified by Vritra exhibited coherent sequence relationships. Sequences from the Vritra- and InterPro-derived datasets were pooled and organized into networks based on >70% pairwise sequence identity. For a well-studied gene such as *frc* (Fig. 2b; additional genes in Supplementary Fig. S1), Vritra-derived sequences showed extensive connectivity with InterPro-derived sequences, consistent with the established functional annotation. In contrast, for under-annotated genes with substantial expansion, such as *xdhA* and *hyuA* (Fig. 2c–d), most sequences formed internally coherent clusters, including groups that were not directly connected to InterPro-derived sequences. These patterns provide sequence-level support for the coherence of the expanded reference space and indicate that Vritra-derived sequences are organized into structured homologous groups rather than being uniformly distributed across unrelated sequence space. Together, these analyses show that Vritra preserves sequence representations supported by existing annotations while extending coverage into sequence space that is underrepresented in current annotation resources.

#### Impact of reference expansion on target-gene read recovery

We next examined whether the expanded sequence space provided by Vritra translated into increased recovery of target-gene reads in real microbiome datasets. We applied the Vritra- and InterPro-derived references independently to the US-men dataset, comprising gut microbiome samples from 308 participants in a subcohort of men enrolled in the Health Professionals Follow-up Study^31^. Raw sequencing reads were aligned to each reference using BLAST separately. We quantified high-quality alignments, defined as reads with >80% alignment coverage and >90% sequence identity, and compared read recovery and sequence identity between the two references. In MGX samples, Vritra recovered a comparable or greater number of target-gene reads than the InterPro-derived reference across the evaluated genes (Wilcoxon signed-rank test, all *P* <0.001; Fig. 3a). The increase was particularly pronounced for less-annotated genes such as *xdhA* and *hyuA*, for which Vritra recovered up to threefold more reads, consistent with their greater sequence representation in the Vritra-derived references (Fig. 2a). Importantly, this increase in read recovery did not compromise alignment quality: mean identities remained ≥97.5% across all genes, and for most genes were significantly higher with Vritra than with InterPro (Supplementary Fig. S2a), with only *ssnA* and *xdhA* showing marginally lower values. These trends were replicated in MTX samples (Fig. 3b and Supplementary Fig. S2b), further confirming that Vritra expands gene-level read recovery without loss of alignment accuracy. The benefit was most pronounced for genes that are poorly represented by existing annotation resources.

**Fig. 3.**
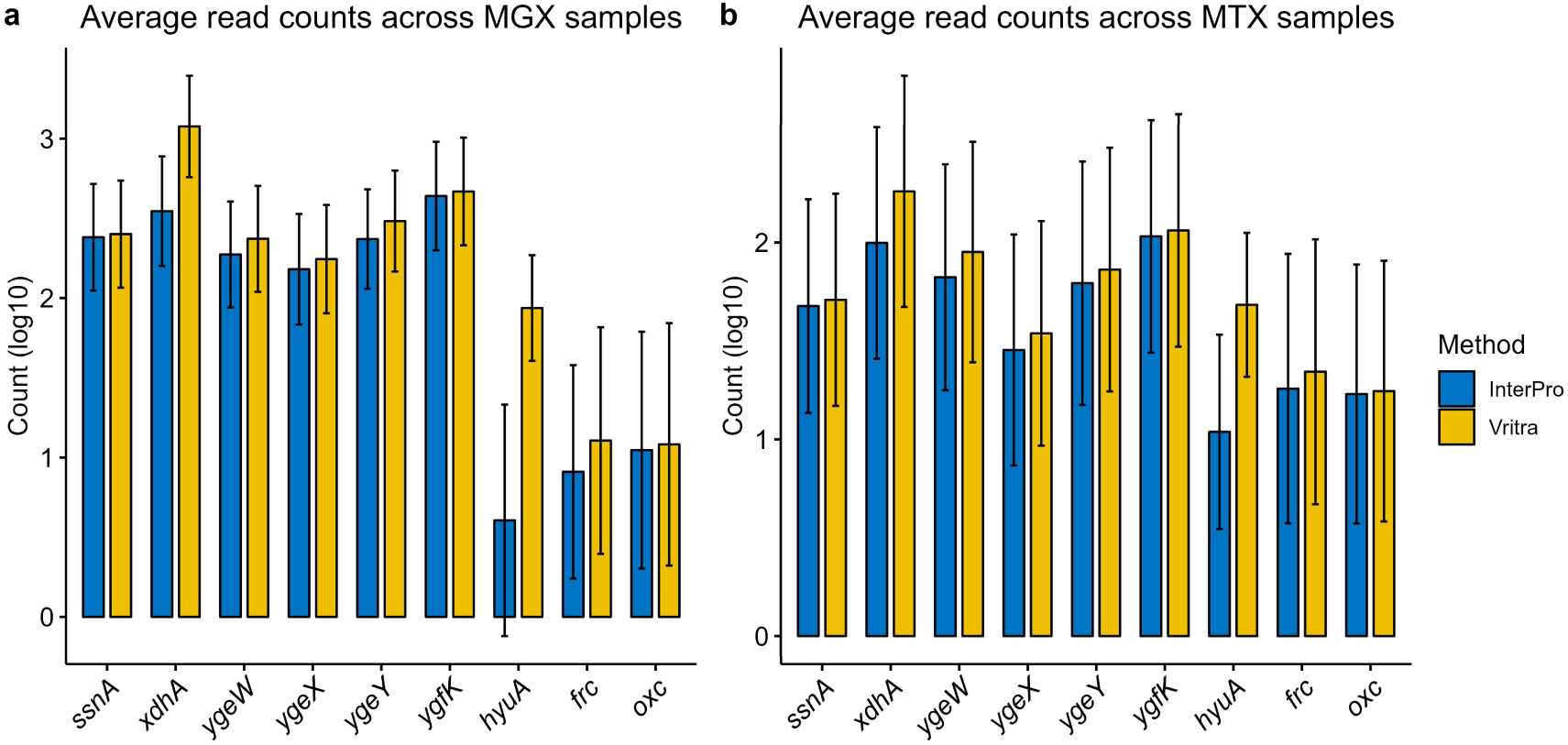
Log10-transformed number of raw sequencing reads aligned to Vritra-constructed and InterPro databases in two gene sets from the US-men cohort. **a**, MGX samples. **b**, MTX samples. Bars represent the mean log10-transformed number of aligned reads, with error bars indicating standard deviation. Statistical significance was assessed using the Wilcoxon signed-rank test for paired samples (all *P* <0.001).

### Downstream analyses with results of Vritra

To illustrate the analytical capabilities of **Vritra**, we examined the association between kidney stone formation and the microbial genes *frc* and *oxc*. Using Vritra’s outputs, we compared genus-level relative abundances between stone formers (SFs, n=58) and non-stone formers (NSFs, n=250) for *frc* (Supplementary Fig. S3) and *oxc* (Supplementary Fig. S4). In MGX samples, no significant differences were observed between SFs and NSFs across the three α-diversity indices (Observed, Shannon, and Simpson) for both genes (Supplementary Fig. S5a, c). In contrast, in MTX samples, the Simpson index, but not the Observed or Shannon indices, was significantly higher in SFs than in NSFs for both *frc* (*P*=0.022; Supplementary Fig. S5b) and *oxc* (*P*=0.0035; Supplementary Fig. S5d). Because the Simpson index is more sensitive to dominant taxa, this result suggests that the microbial communities expressing *frc* and *oxc* are less even and more dominated by specific species in NSFs than in SFs.

Prevalence analyses further revealed modest but consistent shifts: *frc* and *oxc* were more frequently detected in SFs within MGX data, whereas the opposite pattern was observed in MTX data (Supplementary Fig. S6). Specifically, the prevalence of *frc* (*P*=0.054) and *oxc* (*P*=0.041) was lower in SFs than in NSFs in MTX samples. Likewise, RPKM values of *frc* and *oxc* did not differ significantly between SFs and NSFs in MGX data (Supplementary Fig. S7), but were significantly higher in NSFs than in SFs in MTX data for both *frc* (*P*=0.0055; Supplementary Fig. S7a) and *oxc* (*P*=0.022; Supplementary Fig. S7b).

To further examine the contributions of individual species, RPKM and prevalence were decomposed to the species level (*frc* in Fig. 4, *oxc* in Supplementary Fig. S8). Most species identified here overlapped with those reported in our prior oxalobiome study^1^. Notably, in addition to the well-characterized *Oxalobacter formigenes*, we also detected three closely related species (Fig. 4a): *Oxalobacter paraformigenes*, *Oxalobacter paeniformigenes*, and *Oxalobacter aliiformigenes*, which were recently defined^32^ and not captured by the previous pipeline. Consistent with Liu’s results^1^, in both SF and NSF groups, we also observed the shift of high abundance/prevalence of species from genus like *Escherichia*, *Bifidobacterium*, and *Staphylococcus* in MGX data to genus *Oxalobacter* in MTX data (Fig. 4 a, b). To evaluate differences in oxalate-degrading enzyme (ODE) contributions between SF and NSF groups, we quantified the population-level impact of individual genera using Liu’s method^1^, which aggregates genus-specific contributions across samples. In MGX data, the proportion of *Oxalobacter* contributions decreased from 24.1% in NSF to 18.2% in SF (*P*=0.0084), accompanied by a decline in its rank from first to third (Fig. 4 c). A similar trend was observed in MTX data (Fig. 4 d), where *Oxalobacter* contributions decreased from 83.5% in NSFs to 73.3% in SFs (*P*=0.026). Despite this consistent decline from NSFs to SFs across both datasets, *Oxalobacter* remained the dominant genus in MTX samples for both groups. Collectively, these results indicated that oxalobiome activity is more accurately reflected by MTX data, because non-*Oxalobacter* species carrying *frc* and *oxc* showed low-level or no transcription of these genes.

**Fig. 4.**
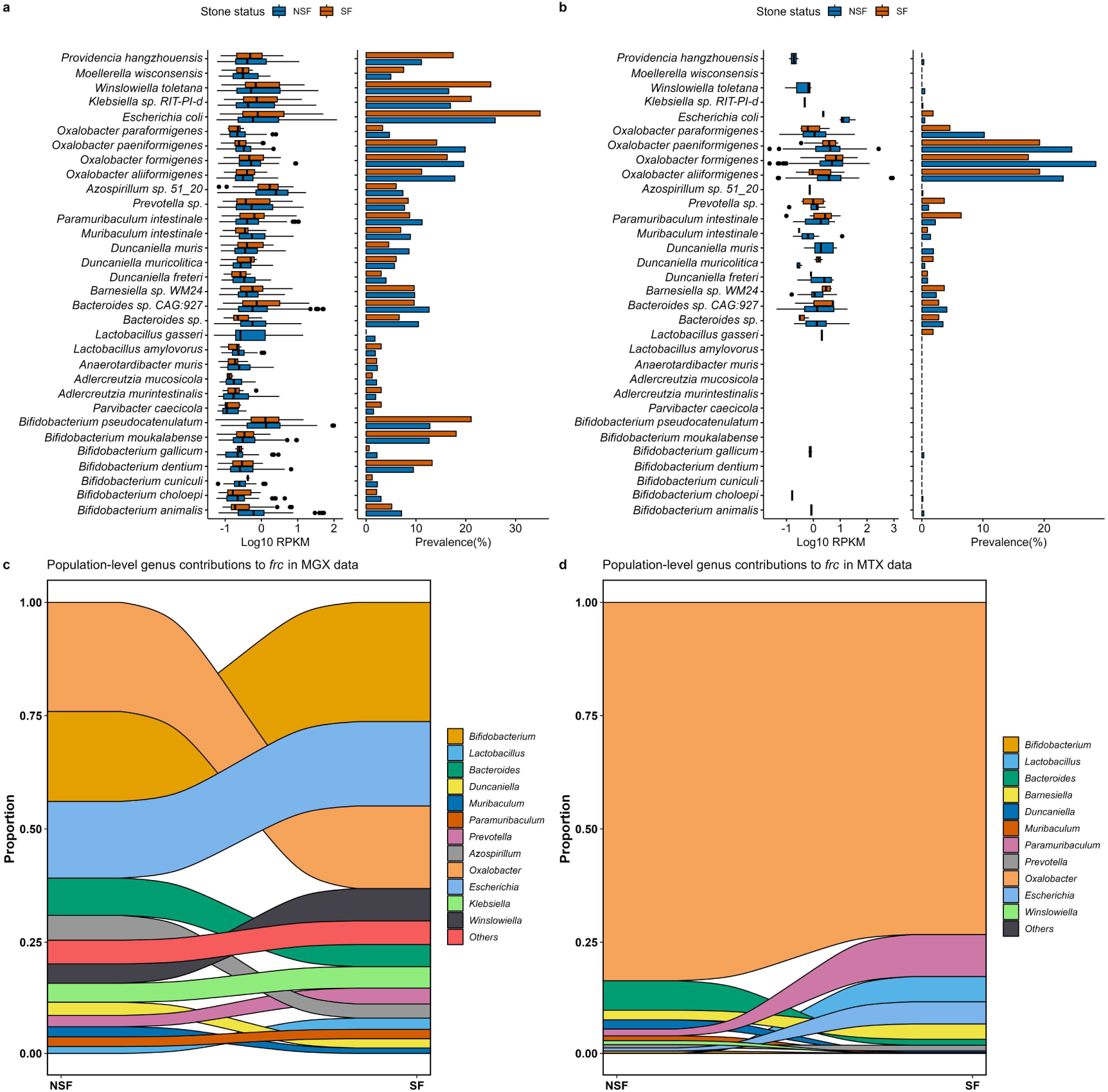
Microbial contributions to *frc* differences between SF and NSF in MGX and MTX samples from the US-men cohort. **a–b**, Abundance and prevalence of species-level *frc*. Box plots show the abundance of microbial *frc* (log₁₀ RPKM) among subjects in whom *frc* is detected, and bar plots show the prevalence (% annotated). **a**, MGX samples. **b**, MTX samples. **c–d**, Population-level contributions of individual genus to *frc* differences between SFs and NSFs, calculated on a relative scale (see Methods). **c**, MGX samples. **d**, MTX samples.

For the urate-degradation gene set, we first quantified the proportional contributions of different genera to the seven genes, following the same approach used for ODE contribution analysis. Unlike the ODE pathway, which was largely dominated by a single genus in MGX data, no genus dominated any of the seven urate-degradation genes in either MGX or MTX samples (Fig. 5a, Supplementary Fig. S9). Moreover, no genus exhibited a marked increase in dominance from MGX to MTX data, as *Oxalobacter* did in the ODE analysis, suggesting that urate degradation is mediated by a broader and more diverse set of microbial taxa.

**Fig. 5.**
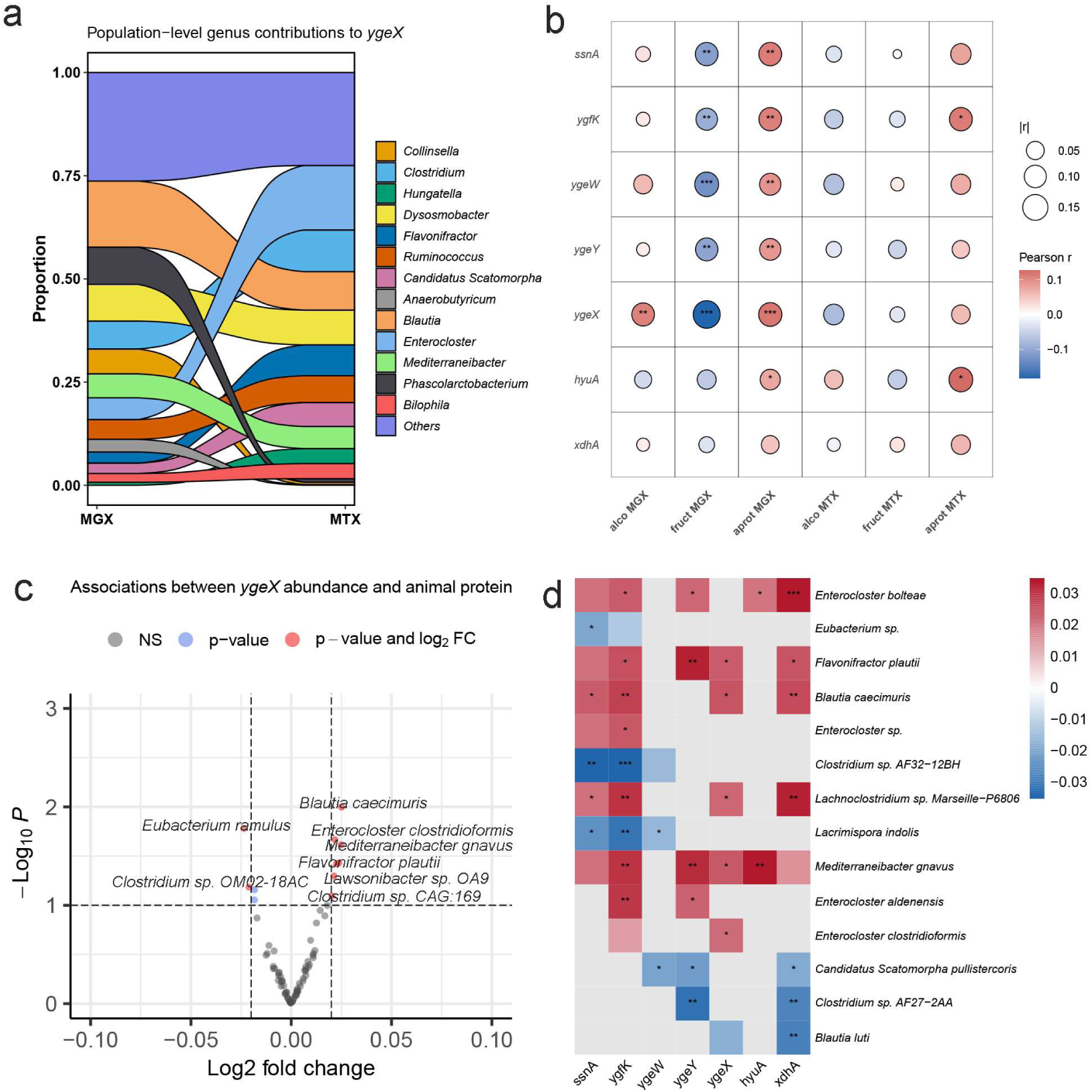
**a**, Population-level contribution of individual genus to *ygeX* differences between MGX and MTX, calculated on a relative scale (see Methods). **b**, Correlations between gene abundance (RPKM) for seven urate-degradation genes and three dietary factors (alco: alcohol; fruct: fructose; aprot: animal protein) in MGX and MTX. **c**, Association between species-level *ygeX* abundance and animal-protein intake; the x-axis indicates the log₂ fold change in gene abundance per unit increase in animal-protein intake. **d**, Associations between species-level gene abundance and animal-protein intake for seven urate-degradation genes; color denotes the log₂ fold change in gene abundance per unit increase in animal-protein intake. Colored blocks without symbols indicate *P* <0.1; * for *P* <0.05; ** for *P*<0.01; *** for *P* <0.001.

We further assessed associations between gene abundances and dietary factors known to influence purine-uric acid metabolism, including alcohol, fructose, and animal protein intake. In MGX samples, significant associations were primarily observed between fructose or animal protein intake and urate-degradation gene abundances, whereas in MTX samples, associations were limited to animal protein intake and the expression of *ygfK* and *hyuA* (Fig. 5b). While gene abundances correlated positively with animal protein, they were negatively correlated with fructose intake, which may reflect dietary confounding: individuals with higher animal protein consumption may have lower fruit intake^33^, consistent with the observed negative correlation between animal protein and fructose in this dataset (Pearson’s *r*=−0.31, *P*<0.001; Supplementary Fig. S10). Mechanistically, fructose increases serum uric acid via hepatic purine degradation^34, 35^, whereas animal protein directly increases exogenous purine load in the gut^36^, explaining why fecal microbiome gene abundances correlated more strongly with animal protein.

To further evaluate species-level associations with animal protein, we performed the differential abundance analysis using ANCOM-BC2^37^. The strongest correlation was observed for *ygeX* in metagenomic samples (Fig. 5b), and corresponding log2 fold changes in species abundance per unit increase in animal protein intake are shown in Fig. 5c, with results for other genes in Supplementary Fig. S11. Notably, *Enterocloster clostridioformis* and *Mediterraneibacter gnavus* (formerly *Ruminococcus gnavus*), identified among species with the largest positive fold changes, have been reported to anaerobically metabolize uric acid^7^, suggesting a role in intestinal urate degradation. Applying filtering criteria (log2 fold change > 0.01 per unit animal protein, p < 0.1, retained if observed for ≥2 genes), we observed consistent directions of fold change across all genes and species (Fig. 5d). Additional species within the genus *Enterocloster* also showed positive associations with dietary animal protein. Although *Blautia caecimuris* has no reported direct role in urate degradation, a related strain (*Blautia* sp. KLE 1732) has been shown to metabolize uric acid in culture^7^.

We also constructed reference databases for *baiB* and *baiE*, two genes that lie at the center of the bile acid 7α-dehydroxylation pathway, and applied them to the IBD Multi-omics Database (IBDMDB)^38, 39^. As shown in Supplementary Fig. S12, Vritra successfully detects the abundance of this under-studied bai gene set, which is poorly represented in InterPro, and identifies the contributing species. Most of these species, including the well-known *Clostridia* taxa *Clostridium scindens* and *Clostridium hylemonae*, have been reported in previous studies to be involved in the bile acid pathway^40^.

## Discussion

High-resolution characterization of specific microbial functional genes and their contributing species remain challenging in MGX and MTX data. Existing frameworks such as HUMAnN enable broad functional profiling of microbial communities, but are not designed to systematically define and resolve user-specified genes at species level. ShortBRED^25^ clusters protein sequences into families based on highly specific marker peptides, but requires users to provide sequences for database construction and may fail to detect reads from short genes (e.g., *oxc* in our previous study^1^).

To address this complementary analytical need, Vritra constructs gene-specific reference databases for user-specified target genes and links them to standardized microbial taxonomy. By integrating sequence-network connectivity with functional annotations, Vritra expands and refines the UniRef-based sequence space to define a core set of functionally relevant sequences, while retaining excluded homologs as decoys to reduce ambiguity during read assignment. The refined sequence space is subsequently linked to UniRef100 and standardized taxonomy to generate compact references for MGX and MTX profiling.

Across oxalate-, urate-, and bile-acid-related genes, Vritra recovered sequence representations supported by established annotations while substantially expanding coverage for less-annotated genes. The expanded sequence sets showed coherent sequence-level network connectivity and increased recovery of target-gene reads in both MGX and MTX data, with the greatest gains observed for poorly annotated genes such as *xdhA* and *hyuA*. For the poorly represented *baiB* and *baiE* genes, Vritra enabled their profiling in real microbiome data despite limited InterPro coverage.

Application to the US-men and HMP2 datasets demonstrated the biological utility of this species-resolved framework. Vritra identified *Oxalobacter* as the major contributor to oxalate-degradation activity and resolved additional recently defined *Oxalobacter* species, while *frc* and *oxc* showed higher relative abundance in NSFs than in SFs. In contrast, urate-degradation genes were distributed across a broader range of taxa, and their abundance was positively associated with dietary animal protein intake. Several urate-degrading species identified by Vritra also showed positive associations with animal protein consumption, including two supported by experimental evidence. Together, these results illustrate how resolving individual functional genes to their contributing species can connect microbial metabolic functions with ecological, dietary, and disease-related contexts.

The current framework is primarily guided by sequence similarity, functional annotations, and standardized taxonomic resources, providing a transparent and reproducible approach that can evolve with improvements in these resources. Future extensions could incorporate additional sequence, structural, or machine-learning-based information to further refine functional discrimination, particularly for poorly characterized genes. More broadly, Vritra provides a flexible framework for translating established and continuously expanding sequence resources into gene-specific references and for resolving the microbial taxa contributing to functions of biological interest.

## Methods

### Construction of the raw UniRef90 dataset

The seed sequence must be supplied by the user and can originate from experimental data, a reference genome, or curated UniProtKB/Swiss-Prot entries on the UniProt^11^ website. In this study, all seed sequences were drawn from UniProtKB/Swiss-Prot and are listed in Supplementary Table S1. The UniRef90 database (version 2025_04) is retrieved from UniProt using the “pre_construct” command provided in the Vritra package. The seed sequences are aligned against the database using BLAST^41, 42^, also implemented in Vritra. Default BLAST parameters are used to allow relatively permissive matches, facilitating the inclusion of a broad set of candidate sequences during the initial stage.

### Build the core UniRef90 dataset via LPA for well-annotated genes

A core UniRef90 dataset can be generated by extending a user-specified target set of pre-labeled sequences, acquired from UniProt or InterPro by searching for the relevant gene or protein names. UniProt, which provides broad sequence coverage and general functional annotations, is recommended in most cases. InterPro, which specializes in the classification of protein families and domains, is preferred when the target gene is well characterized and associated with distinct domain annotations.

The LPA is a graph-based clustering approach in which each node iteratively adopts the most frequent label among its neighbors. In Vritra (Fig. 1a, Step 2), the pre-labeled sequences serve as initial nodes, and labels propagate through the sequence-similarity network. Edges between nodes are defined by pairwise sequence identity, and neighboring nodes are those within a specified similarity radius (70% identity by default). The remaining sequences that do not receive propagated labels constitute the decoy dataset. In the real data analyses in this study, we applied the most stringent rule, allowing each sequence to propagate only to its single nearest neighbor as measured by sequence identity (i.e., identity to the nearest pre-labeled sequence > identity to the nearest decoy sequence). We selected a 70% threshold because large-scale evaluations have shown that clustering performance at 70% retains sensitivity comparable to 90% on massive protein datasets^43^, and more recent adaptive-threshold approaches recommend a lower bound of ∼70% to maintain intra-cluster coherence while reducing redundancy^44^.

### Build the core UniRef90 dataset via iterative search for genes with limited annotation

For genes lacking reliable annotations, Vritra heuristically construct the core UniRef90 dataset from a single seed sequence through iterative search within the initial raw dataset. This procedure parallels Step 2 (Fig. 1a), except that all sequences falling within the search radius are newly assigned the seed label. The process is then iteratively applied to each newly labeled sequence until convergence or until the user-specified maximum number of iterations is reached.

### Re-cluster UniRef100s into species

To improve species-level representativeness and downstream analysis accuracy, we re-clustered UniRef100 protein sequences based on the relationship between UniRef90 clusters and species. These relationships were categorized into four types, and representative sequences were selected accordingly (Fig 1a, Step 3):

1. **Single UniRef90–Single Species**: A UniRef90 contains all UniRef100 sequences from a single species, and no sequences from other species. The representative sequence is served by the centroid sequence of the UniRef100s within the UniRef90—that is, the sequence with the highest average identity to all others in the set. Let *S* = {*S*_1_, *S*_2_,…, *S_n_*} be the set of UniRef100 protein sequences within a UniRef90 cluster. Let *Iden_i,j_* denote the pairwise sequence identity between sequences *S_i_* and *S_j_*. The centroid sequence *S_c_* ∈ *S* is defined as equation (1):

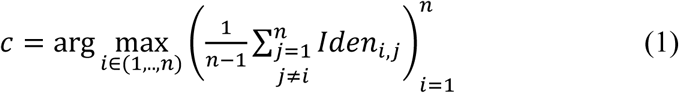
2. **Multiple UniRef90–Single Species**: Multiple UniRef90s collectively contain all UniRef100 sequences of a single species, and each UniRef90 in this group only includes sequences from that species. As in the first category, a centroid sequence is selected from each UniRef90 cluster based on average pairwise identity and retained for downstream database construction. However, if the centroid sequences from different UniRef90s share high similarity (pairwise identity > 96%, red circle in the second category in Fig 1a, Step3), their corresponding UniRef100 sets are merged, and a new centroid is computed from the pooled sequences. In other words, highly similar UniRef90s are consolidated to produce a single, more representative centroid sequence. Notably, the 96% identity threshold is consistent with the approximate average nucleotide identity (ANI) boundary used to delineate species in the Genome Taxonomy Database (GTDB). This threshold also corresponds to roughly one amino acid mismatch per 33 residues (or ∼99 base pairs), which aligns with the minimum read length of most next-generation sequencing (NGS) platforms.
3. **Single UniRef90–Multiple Species**: In this scenario, a UniRef90 contains sequences from multiple species, and all sequences from each species are found exclusively within this UniRef90. To resolve species-level representation, all UniRef100 sequences within the UniRef90 are re-clustered using the Affinity Propagation algorithm^45^ (implemented via the scikit-learn library^46^), based on pairwise sequence identity. A centroid sequence is then selected for each resulting cluster. If all sequences within a cluster originate from the same species, the centroid is labeled accordingly. For mixed-species clusters, the centroid is labeled with the species that contributes the majority of sequences, while full taxonomic composition is recorded in the output for reference.
4. **Multiple UniRef90–Multiple Species**: In this category, several UniRef90 clusters contain UniRef100 sequences from multiple species, and sequences from the same species may appear in more than one UniRef90. We first examine whether sequences from the same species with high identity (>96%) are erroneously split across different UniRef90s. If so, the corresponding UniRef100 sequences will be merged, and a new centroid will be computed from the combined set (as illustrated by the red circle in the fourth category of Fig. 1a, Step 3). Otherwise, the corresponding UniRef100 sequences from different UniRef90s will not be merged. After resolving these cases, the remaining UniRef100 sequences are reassigned to either category-1 or category-3, depending on their specific composition.

This strategy ensures that the selected representative sequences are both taxonomically coherent and structurally representative of the full set of protein sequences for each species.

### Microbiome Differential Abundance Analysis

Differential abundance results were filtered using the pseudo-sensitivity diagnostic from ANCOM-BC2, retaining only taxa with absolute pseudo-sensitivity values for the covariate aprot10a less than 0.1 to ensure robustness of the estimated log-fold changes.

## Author contributions

Conceptualization: B.Z., H.L.

Data was obtained by: B.Z., G.C., L.N., H.L. Bioinformatic method development: B.Z., M.L., L.N., H.L.

Real data analysis and biological interpretation: B.Z., M.L., G.C., G.P., H.C., L.N., H.L. Programming and Software: B.Z.

Writing-Original Draft Preparation: B.Z., H.L.

Writing-Review and editing: B.Z., M.L., G.C., G.P., H.C., L.N., H.L.

## Supplementary material

Supplementary material is available at *mSystems* online.

## Conflict of interest

None declared.

## Funding

This work was supported by the National Institutes of Health [R01LM014085 to H.L., B.Z.; R01DK137473 to H.L., B.Z., G.C., L.N.; R01DK129675 to H.L., B.Z., L.N.; 1R01DK141277 to H.L., B.Z., G.P.].

## Data availability

The raw sequencing data from the US-men cohort is available on NCBI BioProject (https://www.ncbi.nlm.nih.gov/bioproject/354235) with accession number PRJNA354235. Metagenomic sequencing data from the IBDMDB can be accessed through the database portal (https://ibdmdb.org/results).

## Code availability

Vritra was implemented in Python 3 on a Linux platform and is freely accessible at the project repository with detailed tutorials (https://github.com/BoyanZhou/Vritra) under the MIT license. The repository also contains the scripts used to generate all figures in this study.

